# A unified smoothing framework for protein domain bigram model

**DOI:** 10.64898/2026.06.14.732219

**Authors:** Xiaoyue Cui, Gautam Iyer, Dannie Durand

## Abstract

**Motivation:** Biomolecular sequences can be represented as strings over an alphabet, an analogy that has motivated many applications of computational linguistic techniques to biological problems. However, such methods must be adapted to the characteristic scale and organization of biomolecular data. Here, we consider the problem of bigram smoothing for multidomain protein architectures, where domain bigram frequency data is extremely sparse and differs from textual data in alphabet size, string length distribution, the relationship between bigram and unigram frequencies, tandem repeat lengths, and the distribution of domain adjacencies. Moreover, some domain combinations are unobserved because they are biologically incompatible, others because the data are incomplete. A smoothing method that distinguishes these two cases is required.

**Results:** We propose a unified smoothing framework based on interpolation that can be tuned to accommodate different bigram data characteristics. Within this framework, we design specific model variants suited to protein domain bigram data: these assign low adjusted counts to pairs that are likely incompatible, while making appropriate adjustments for undersampled pairs. We demonstrate empirically that this approach distinguishes the two cases while preserving the characteristic signatures of multidomain data.

**Availability and implementation:** Implementations of smoothing methods, the scripts used to generate all results presented in this paper, and the curated lists of extracellular and DNA-binding domains are available at https://codeberg.org/xcui297/protein-domain-smoothing.

## Introduction

A multidomain protein is a mosaic of sequence segments that encode *domains*, structural or functional modules that are found in diverse, otherwise unrelated, contexts. The ability of a domain to take on its characteristic fold is independent of the surrounding sequence. The *domain architecture* of a protein, that is, its domains in N-to C-terminal order, can be determined by scanning a database of probabilistic domain models, e.g. [7, 28, 35, 36]. This domain architecture representation is a widely used abstraction for investigating the evolution of the protein domain repertoire [21, 33, 40, 11, 14], plasticity in domain order [4, 24, 37], domain occurrence graphs [34, 22, 10, 30], and domain promiscuity, i.e., the propensity of a domain to co-occur with many other domains [26, 6, 5, 9, 10, 38].

The matrix of ordered domain pair or *bigram* frequencies is a fundamental data structure used in many of these applications. It provides a characteristic signature of the genome’s protein function repertoire [41], as well as an implicit representation of Nature’s “design principles” for multidomain proteins. The bigram frequency matrix can be used to estimate transition probabilities in a Markov model of domain architecture evolution [12] and domain architecture probabilities using a first order approximation consisting of the product of conditional bigram probabilities (Equation (2)).

Bigram frequencies are estimated from domain pair counts observed in empirical domain architecture data. Unfortunately, domain architecture data is sparse and noisy, which poses both practical and fundamental challenges to downstream analyses. Domain pairs that do not appear in any domain architecture in the input data (*unseen pairs*) have zero-valued bigram counts, which cause mathematical problems. In addition, the counts of domain bigrams that are observed at least once (*seen pairs*) may be inaccurate due to various sources of noise and missing data. Some bigrams are underrepresented due to errors in sequencing and in gene and domain annotation. Domain combinations that are encoded by genomes that have yet to be sequenced are also not accounted for. Still other combinations may not be encoded in any genome currently on the planet because evolution has not yet “discovered” all functionally viable domain architectures. All of these can contribute to uncertainty in both seen and unseen pair frequencies.

Interpretation of unseen bigram frequencies is further complicated because not all zero-valued entries are due to data incompleteness. The domain combinations that occur in nature are highly constrained by biological forces. Some domain pairs may not be observed because they are genetically, structurally, or functionally incompatible. For example, the beta-beta-alpha zinc finger (Znf) domain (SF 57667) and the immunoglobulin domain (SF 48726), two of the five most common domains in human domain architectures according to the Superfamily database [28], never co-occur in the same protein. The Znf domain is a DNA-binding domain found in transcription factors that localize to the nucleus. In contrast, immunoglobulin (Ig) domains are primarily found in the extracellular regions of proteins with roles in cell adhesion and allorecognition [20], suggesting that these domains do not co-occur because they are associated with incompatible cellular locations. Our confidence that Znf and Ig domains truly do not co-occur is boosted by the fact that both domains are highly abundant when considered individually.

Sparse and noisy data can be mitigated by smoothing techniques that eliminate zero-valued entries and make adjustments to compensate for observation errors. The smoothing protocol should be consistent with the underlying properties of the data specific to the application at hand. To accomplish this, a smoothing method should be both tunable and non-uniform. That is, it should be parameterized to allow the overall amount of adjustment to be tuned to account for the degree of noise in a particular data set. Further, it should permit different amounts of adjustment to different data entries to reflect the fact that some entries are more reliable than others. Finally, the adjustments should be small enough to preserve the characteristic structure of the data.

A smoothing method for domain architecture data must seek to balance the impact of incompleteness, on the one hand, and biological constraints, on the other. It should assign non-zero counts to unseen pairs, taking into consideration the fact that some pairs may be unseen due to data incompleteness, while others are truly absent. It should further adjust the counts of seen pairs to account for incompleteness. Prior multidomain analyses have relied on smoothing techniques originally developed in the field of natural language processing (NLP), although language data has very different underlying properties. Cui et al. [12] used additive smoothing, where the raw data are adjusted by adding a constant *pseudocount* to the number of observed pairs for every bigram. With additive smoothing, the degree of smoothing can be tuned by modifying the pseudocount, but is applied uniformly to all data points. Good-Turing smoothing [17], which was used by Yu et al. [41], assigns the same adjusted count to all bigrams with the same cardinality. While Good-Turing is non-uniform, it is not tunable; rather, all entries with the same cardinality are adjusted in the same way. Both additive and Good-Turing smoothing treat all unseen pairs identically. For protein domain bigram data, this is problematic because these methods cannot account for the possibility that some unseen pairs are not observed because they are incompatible, while others are simply due to missing data.

### Our contributions

We introduce a general bigram smoothing framework that is both tunable and non-uniform and that easily generalizes to higher order *n*-gram models. We then present a realization of this model that is tailored specifically for protein domain bigram smoothing. We restrict our attention to bigram models; higher order *n*-gram models are not viable for domain architectures, even for *n* = 3, as domain architecture data are extremely sparse.

Our model is informed by the Znf-Ig example, in which two domains that both occur frequently have never been observed to co-occur. Generalizing from this example, we base our method on the assumption that single domain (*unigram*) frequencies provide information about the reliability of the observed domain pair data: pairwise counts of more frequently occurring domains should be regarded as more reliable. Following this rationale, we design our smoothing strategy to account for data in a way that makes make smaller adjustments to portions of the data that represent more reliable observations. This is achieved by a parameterized model that interpolates between the observed counts and a target count distribution. We explore several variants of this framework with different parameterizations and target distributions.

We evaluate smoothing performance empirically on human domain architecture data (Figure 1). We demonstrate that adjustments to bigram counts do indeed decrease as the associated unigram frequencies increase and verify that our smoothing method assigns lower scores, on average, to a curated set of incompatible pairs with high unigram frequencies, compared with unseen bigrams in general. We further compare the performance of the proposed smoothing variants using figures of merit designed to assess how well smoothing completes incomplete data.

**Fig. 1.**
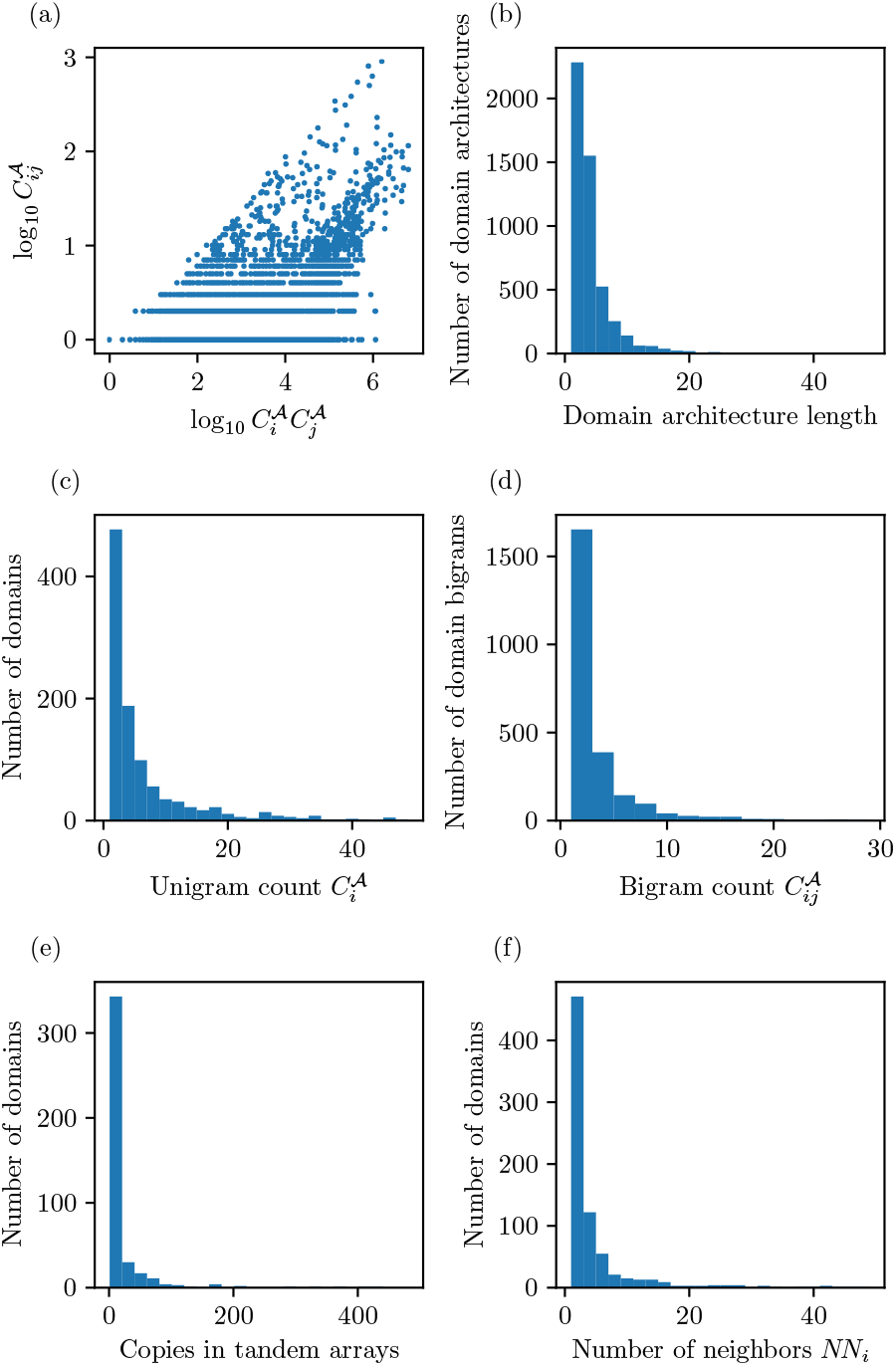
Characteristic properties of human domain architecture data. (a) The relationship between product of unigram counts and bigram count. (b) Distribution of domain architecture lengths. Distributions of (c) unigram counts, (d) bigram counts, (e) the number of copies found in tandem arrays, and (f) the number of distinct direct neighbors, i.e., domain promiscuity. All histograms are plotted over a restricted range for clarity. Full scale distributions are shown in Figure S1.

Finally, we demonstrate that our smoothing method preserves characteristic features of domain architecture data, including domain repetitiveness, promiscuity (the propensity of domains to co-occur with many other domains) and order constraints. Domain order constraints, in particular, are a challenge for smoothing domain architecture data, because the data is too sparse to model n-gram statistics accurately, for *n >* 2. However, our empirical analysis of genome-wide human bi- and trigram statistics shows that bigram frequencies implicitly encode longer range order constraints. Further, in case studies of selected signaling proteins, calculations based on smoothed bigram statistics are able to distinguish genuine domain architectures from permutations of the same domains.

## Methods

We first introduce notation used in subsequent sections, describe the domain architecture data used for empirical analyses, and set up a framework for performance comparisons across proposed smoothing variants.

### Notation

Let *D* = {*D*_1_, *D*_2_,…, *D*_*N*_} be the set of all protein domain families in genuine data. This alphabet is augmented with two additional tokens, *D*_*S*_ and *D*_*E*_, that indicate the start and end of an architecture. A domain architecture *A* of length *n* is a finite sequence of domains 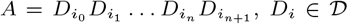, where 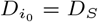 and 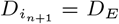. Given a set of unique domain architectures, 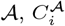 and 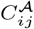 represent the number of instances of domain *D*_*i*_, i.e., the unigram count of the domain, and the number of ordered pairs *D*_*i*_*D*_*j*_, i.e, domain bigram count, respectively. The total bigram count is denoted *C*_0_ = ∑*i,j C*_*ij*_.

The probability of a domain architecture can be computed using the chain rule of probability:

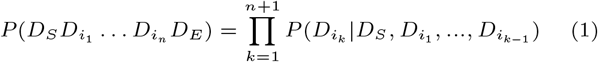

A *bigram model* provides a first-order approximation of this probability, wherein each term corresponds to the conditional probability of a observing domain *D*_*i*_ given the preceding domain, *D*_*i*− 1_:

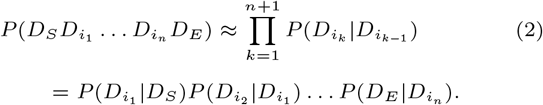

The conditional probability of observing *D*_*j*_ following *D*_*i*_ can be estimated with the maximum likelihood estimator (MLE), which we denote simply by

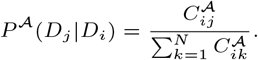

The MLEs of the domain unigram and bigram probabilities can also be defined in terms of unigram and bigram counts, yielding

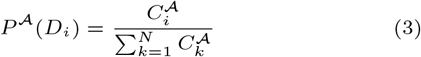

and

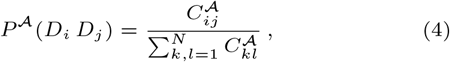

respectively.

The quantity 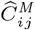 denotes the adjusted count for bigram *D*_*i*_*D*_*j*_ for a given smoothing method, *M*. Additive smoothing, the method used by Cui et al. [12], addresses unseen data by adding a pseudocount *Ψ >* 0 to the count of all bigrams, yielding

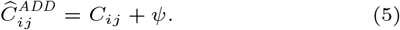

Since additive smoothing is both widely used and extremely simple, we use it as a baseline for comparison with the smoothing strategies introduced here.

Good-Turing smoothing, which has also been used in prior domain bigram studies [41], is calculated as follows: Let the *cardinality count*, denoted *N*_*c*_, be the number of distinct bigrams with observed count *c*, that is

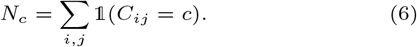

Smoothing is performed by adjusting the cardinality counts, *N*_*c*_,*c* = 0, 1,…, which are then used to obtain adjusted bigram counts. The adjusted cardinality count, 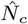, is determined by a linear regression fit to log *Z*_*c*_ versus log *c*, where

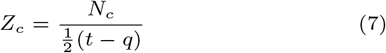

and *q, c, t* are the subscripts of three consecutive non-zero counts, *N*_*q*_, *N*_*c*_, *N*_*t*_. To accommodate edge cases, we set *q* = 0, when *c* = 1. When *c* = *c*_max_, the greatest nonzero count, the denominator is replaced with *c*_max_ − *q*, resulting in

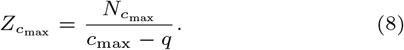

The adjusted cardinality counts are used to obtain adjusted bigram counts. For any seen bigram (*C*_*ij*_ *>* 0), we have

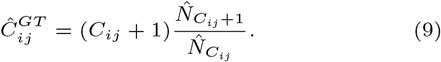

For unseen bigrams (*C*_*ij*_ = 0), 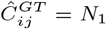.

We can see that this method adjusts all bigrams with the same cardinality in the same way. In particular, all unseen bigrams are assigned the same adjusted count.

### Test data curation

We evaluate the behavior and performance of models proposed in this study using domain architecture data obtained from the online Superfamily database, version 1.75 [19]. Domain annotations of 58,023 human protein sequences were downloaded and parsed using the pre-processing module of DomArchov [12]. The resulting dataset has 5, 031 unique domain architectures, with mean, median, and maximum lengths of 4.2, 3 and 249, respectively. From these, we extracted the domain alphabet and calculated domain unigram and bigram frequencies. The domain alphabet has *N* = 1, 109 distinct superfamilies. A total of 2, 526 distinct domain bigrams are observed, comprising ~0.2% of *N*^2^ ≈ 10^6^ possible bigrams (Figure 1(d)).

Our smoothing method is designed to assign minimal adjustments to unseen pairs with high unigram frequencies. To test whether this goal is achieved, we require a set of unseen bigrams composed of domains with high unigram frequencies that are likely biologically incompatible. The example of the Znf and Ig domains, which are highly abundant but never observed together, suggests a strategy for curating such a set. We posit that DNA-binding (DB) domains, which are predominantly localized to the nucleus, will rarely be observed adjacent to domains that are found exclusively or primarily in extracellular proteins or in the extracellular regions of receptors, so-called extracellular (EC) domains. We curated a set of 99 DNA-binding domains and a set of 35 domains with primarily or exclusively extracellular localization, as described below.

For extracellular domains, we obtained an initial list of candidates from two prior studies [3, 27]. PFAM domains that are described as extracellular by Bányai and Patthy [3] were considered further if they have a corresponding entry in the Superfamily database. Of these, domains that have at least one publication linked on InterPro or Uniprot supporting the extracellular localization of either the domain or a protein containing the domain were retained. In addition, Naba and colleagues [27] provide a list of InterPro domains that define the core complement of extracellular matrix proteins, identified using a bioinformatic pipeline, followed by manual curation. Among these, any domain that has a Superfamily counterpart was also included in our study. The combined set comprises 35 EC domains (Table S2).

For DNA-binding domains, Superfamily domains that are either (a) in the DNA-binding category on Superfamily or (b) described as DNA-binding in the description on the Superfamily site provided an initial set of candidates. Those candidates that have a publication linked on Superfamily, InterPro, or Uniprot supporting their DNA-binding function were retained. We augmented this list using the “confident set” of domain-ligand interactions from InteractDome 0.3 [23], defined to be the set of interactions “with nonidentical instances found across at least 3 structural co-complexes and a cross-validated precision of at least 0.5”. Any PFAM domain that has a DNA-ligand in the confident set was added to our list if it has a counterpart in Superfamily. The entire curation procedure resulted in 99 DNA-binding (DB) domains (Table S3), in total.

Out of the 6,930 possible bigrams formed by selecting one EC and one DB domain, in either order, 5,998 (87%) are unseen in human domain architectures. The unigram counts of both the EC and the DB domains (Figure S6) are stochastically greater than those of domain superfamilies, overall (*p <* 0.0001, *U >* 4E3, one-sided Mann-Whitney U test). The products of unigram counts (*C*_*i*_*C*_*j*_) for EC-DB (or DB-EC) pairs are also stochastically greater than those for domain pairs, in general (*p <* 0.0001, *U* = 3E8, one-sided Mann-Whitney U test).

### Calibration across models

In Results, we introduce tunable smoothing models, where the degree of smoothing exerted by model *M* is controlled by a parameter, *ε*_*M*_. In order to obtain a fair comparison across model variants, we calibrate the values of this parameter using additive smoothing as a reference. For each model, the value of *ε*_*M*_ is selected such that the total change in seen bigram counts due to smoothing with method *M* is the same as the total change obtained by additive smoothing with parameter *Ψ* ∈ {0.0002, 0.0004, 0.0006, 0.0008, 0.0010, 0.0012, 0.0014, 0.0016}. Let

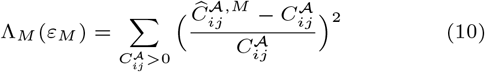

denote the total adjustment resulting from smoothing with model *M*. For each value of *Ψ*, we select a value of *ε*_*M*_ so that

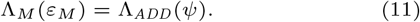

The values of *ε*_*M*_ resulting from this calibration process are given in Table S1. The results presented in Figures 3, 4, S1, S2 and S3 are indexed by this *effective pseudocount*, that is, the value of the pseudocount *Ψ* used to calibrate the models, rather than the actual values of *ε*_*M*_ used in each model.

### Assessing completion of incomplete data

To assess how well the proposed smoothing strategies complete incomplete data, artificial incomplete data sets were constructed by discarding a random sample of *x*% of the genuine architectures, *x* ∈ {1%, 2%,…, 10%}, to obtain reduced sets, denoted *R*. To account for sample variance, 1000 replicates of this process were conducted for each value of *x*.

For each reduced set, *R*, incomplete bigram counts, denoted 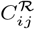, were tabulated from the remaining domain architectures in the set. Unigram frequencies are estimated on the complete set *A* to ensure that all domain superfamilies in the original domain alphabet are retained. Smoothing was applied to the resulting incomplete bigram counts, 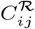, to obtain adjusted bigram counts, denoted by 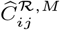, where *M* indicates the smoothing method used.

Three measures were used to assess how well smoothing completes incomplete data:

The *root mean square bigram frequency deviation*, defined by

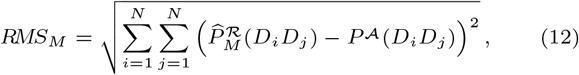

assesses how well the smoothed bigram frequencies derived from the reduced data set approximate the original, genuine bigram frequencies. Here

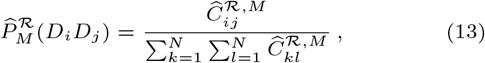

denotes the smoothed incomplete bigram frequency obtained with smoothing method *M*. A smaller value of *RMS*_*M*_ indicates more effective smoothing. If the smoothing procedure perfectly corrected for the incompleteness of the data, this quantity would be zero.

The *architecture probability deviation*, defined by

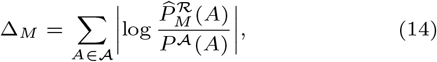

measures how well domain architecture probabilities estimated by applying smoothing model, *M*, to incomplete data approximate the MLE probabilities based on complete genuine data. Here the probability of domain architecture *A* in the numerator of (14) is estimated using the bigram model (2) with conditional probabilities derived from smoothed, incomplete counts:

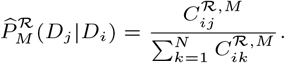

A smaller deviation #_*M*_ indicates a better fit to the domain architecture probability estimated using bigram frequencies derived from the complete set *A*, without smoothing.

Finally, the *log probability of genuine domain architectures*, defined by

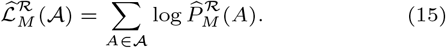

is used to determine which model variant maximizes the probability of the genuine data

## Results

### Customizable Unified Interpolation smoothing

We propose a Customizable Unified Interpolation (CUI) smoothing framework that is tunable and supports flexible and non-uniform adjustment to *n*-gram counts. When *n* = 2, for every bigram *D*_*i*_*D*_*j*_, we obtain an adjusted count 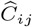 by interpolating between the observed bigram count *C*_*ij*_ and a user-specified target bigram count distribution, 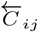. This is achieved by minimizing a *modification cost*,

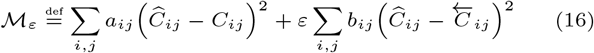

where *ε* ≥ 0 is a parameter that controls the amount of smoothing. The trust parameters *a*_*ij*_, *b*_*ij*_ are chosen to reflect our confidence in the observed pairwise counts *C*_*ij*_ and the target pairwise counts 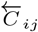, respectively.

The adjusted counts 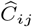 that minimize the cost *M*_*Ψ*_ can be found by equating the partial derivatives 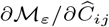 to 0. This gives

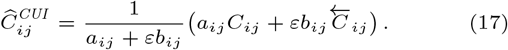

Equation (17) shows that the smoothed bigram distribution is indeed an *interpolation* between the observed bigram distribution and the target distribution. Further, the adjusted bigram counts converge to the actual counts when the smoothing parameter, *ε*, approaches zero:

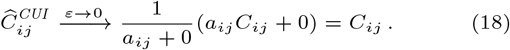

This demonstrates that our smoothing framework makes small adjustments to observed counts when *ε* is small. This observation is not sensitive to the choice of *a*_*ij*_, *b*_*ij*_ and 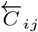.

Equation (17) is a general smoothing framework that can be tailored to suit specific assumptions of many types of bigram data and desired smoothing behaviors, by choosing the target pairwise count distribution 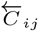, and trust parameters *a*_*ij*_, *b*_*ij*_ based on application-specific considerations. The *larger a*_*ij*_ (or the *smaller b*_*ij*_), the greater the weight given to the observed pairwise counts *C*_*ij*_, and the smaller the weight given to the target pairwise counts 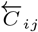.^1^

Note that while we have introduced this framework for the case where *n* = 2, it generalizes easily to larger values of *n*. For example when *n* = 3, the modification cost based on trigrams is

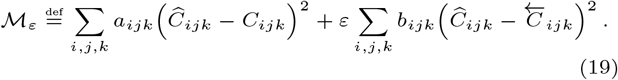

Minimizing (19) gives

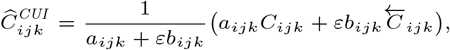

where the 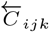 is the target trigram distribution and the weights *a*_*ijk*_, *b*_*ikj*_ are trust parameters as before.

We next present our customization of this generalized smoothing framework for multidomain architecture data. Our goal is to design *a*_*ij*_, *b*_*ij*_, and 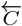, so that the adjusted bigram count distribution has properties that are desirable for modeling multi-domain architectures. We present our specific design and explain how it respects characteristic properties of multidomain architectures. We further discuss why certain alternative designs are less suitable.

### Smoothing protein domain bigram data

Since there is more opportunity to observe more abundant domains, we expect that the observation error for high frequency domains is smaller. Thus we would like a higher trust in observations of such domains and a lower confidence in the target distribution 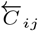.

This can be accomplished by selecting the trust parameters *a*_*ij*_ and *b*_*ij*_ to up-weight the observed counts (*a*_*ij*_ *-weighting*)

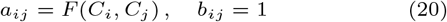

or down-weight the target counts (*b*_*ij*_ *-weighting*)

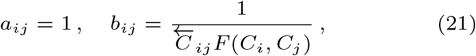

where *F* is a function that increases to infinity as *C*_*i*_, *C*_*j*_ ⟶ #x221E;.

While *a*_*ij*_-weighting (20) may appear more natural at first sight, *b*_*ij*_-weighting (21) has asymptotic properties which are more desirable in our situation. We explain the reason for this below, and will now focus our attention on the *b*_*ij*_-weighting strategy (21).

For *b*_*ij*_-weighting, Equation (17) simplifies to

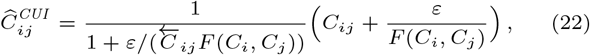

when the weights are chosen as in (21). Two natural choices for the function *F* are

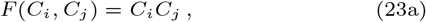

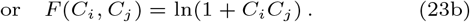

Note that in both cases *F* (*C*_*i*_, *C*_*j*_) increases to infinity as *C*_*i*_, *C*_*j*_ ⟶ ∞. Moreover, both (23a) and (23b) give a numerically stable choice in (22) because the function *F* is bounded away from 0. One advantage of choosing *F* according to (23b), as opposed to (23a), is that human domain architecture data has a characteristic distribution of *C*_*i*_*C*_*j*_ with values ranging between 1 and 10^6^ (Figure 1(b)).

For interpolation, we consider two candidate target bigram count distributions:

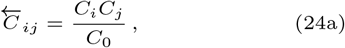

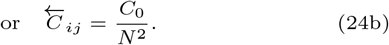

These choices reflect the expected bigram counts under the unigram model and the uniform distribution, respectively. Under the unigram model (24a), the bigram frequencies are given by *P* (*D*_*i*_*D*_*j*_) = *P* (*D*_*i*_)*P* (*D*_*j*_). In other words, the corresponding bigram frequency distribution is such that the marginal distribution exactly coincides with the observed frequency distribution. Interpolation to the uniform distribution of bigrams (24b) makes minimal assumptions and has the potential advantage of robustness, given the noisiness of domain architecture data.

We have proposed two choices of the trust parameter *b*_*ij*_ and two choices of the target distribution, 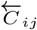, leading to four models overall (Table 1). For all four variants, smoothed bigram counts 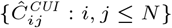 can be calculated in *O*(*N* ^2^) time. We next consider the properties of these variants in light of our design goals.

**Table 1.**
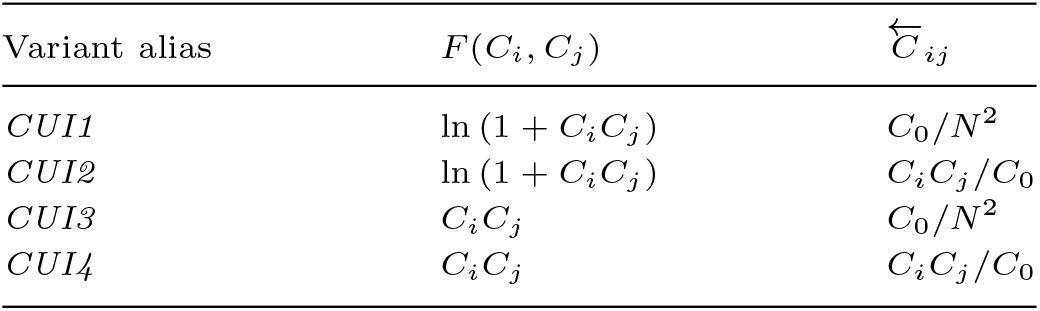
Model variant aliases for method proposed in (21).

### Model interpretation

Our guiding principle is that the observed count of bigrams composed of domains with high unigram frequencies should be treated as more reliable. This can be quantitatively captured by the following properties.

1. The smoothed counts of both seen and unseen pairs should converge to the actual counts as *C*_*i*_, *C*_*j*_ ⟶ ∞.
2. Adjustments to unseen bigram counts should decrease as unigram frequencies increase.

As we illustrate below, both of these expectations are met for *b*_*ij*_-weighting, but not for *a*_*ij*_-weighting. A key design feature in our smoothing scheme is the incorporation of the target count 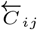 in *b*_*ij*_, which ensures that these two properties hold regardless of the specific choice of 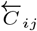.

### Convergence to C_ij_

Our design was guided by the assumption that observed bigram counts are more reliable when the associated unigram counts are large. We seek parameter choices such that 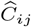 approaches *C*_*ij*_, as unigram counts increase. To understand the adjustment our framework makes in terms of the trust parameters, it is convenient to define the ratio 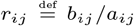, which measures the balance between the confidence in the target distribution and trust in the observed data. Using the fact that when *z* is small,

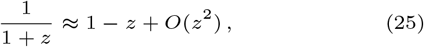

the right hand side of (17) can be approximated by

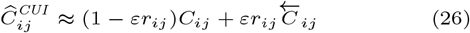

when *εr*_*ij*_ is small. Thus, to make relatively smaller adjustments to certain bigram counts, we need to choose our trust parameters so that 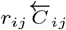 is relatively smaller for these bigrams.

For *b*_*ij*_-weighting (21, 22), we see that the second term in (26) is

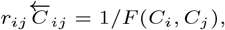

which vanishes as *C*_*i*_, *C*_*j*_ ⟶ ∞. Thus, 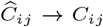 as *C*_*i*_, *C*_*j*_ ⟶ ∞, consistent with our goal of making smaller adjustments to the observed pairwise counts of more frequently occurring domains.

In contrast, with the *a*_*ij*_-weighting (20), the second term in (26) is

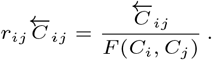

While we still have *r*_*ij*_ = 1*/F* (*C*_*i*_, *C*_*j*_) ⟶ 0 as *C*_*i*_, *C*_*j*_ ⟶ ∞, which is desirable, 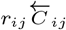 may still be large when *C*_*i*_ and *C*_*j*_ are large. As a result, the second term in (26) may not vanish as *C*_*i*_, *C*_*j*_ ⟶ ∞, and the adjustment to bigram counts of frequently occurring domains may be relatively large. This is not desirable for domain architecture data.

### Unseen bigrams

Adjusting the counts of unseen bigrams is a crucial, but challenging goal of smoothing domain architecture data. Observed data is incomplete in its nature. Pairs that are not observed in the current data may be present after more sampling; alternatively, unseen pairs may be absent because they are selectively disadvantageous. We take advantage of unigram counts to obtain a smoothing function that captures both phenomena. If *D*_*i*_*D*_*j*_ is not observed despite the fact that both domains are common, then it is likely *D*_*i*_*D*_*j*_ is not observed because the pair is biologically incompatible, and so the count should receive a very small adjustment.

For unseen bigrams (*C*_*ij*_ = 0) with *b*_*ij*_-weighting, Equation (22) reduces to

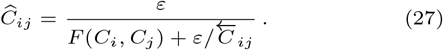

For all variants in Table 1, 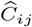 decreases as a function of *C*_*i*_*C*_*j*_, provided that *C*_*i*_*C*_*j*_ ≳ *ε*. That is, when the unigram counts are larger, the adjusted bigram count *decreases* with the unigram counts.

In contrast, with *a*_*ij*_-weighting (20), for unseen bigrams we have

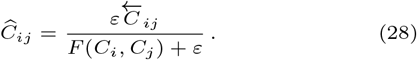

Adjusted bigram counts increase with the unigram counts whenever 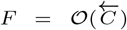, which is not desirable for domain architecture data.

To probe the relationship between 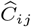 and *C*_*i*_*C*_*j*_ empirically, we applied all four smoothing methods in Table 1 to unseen bigrams in genuine human domain architecture data (see Methods) and compared the results with those obtained with *a*_*ij*_-weighting and the two smoothing methods used in NLP. We see that all variants in Table 1 result in decreased adjusted counts for larger unigram counts, as designed (Figure 2(a)). In contrast, smoothed bigram counts obtained with *a*_*ij*_-weighting display an increasing trend, which is not desirable. We further examine the impact of additive and Good-Turing smoothing methods on unseen bigram counts. Both additive and Good-Turing treat all unseen bigrams identically, resulting in the same value of 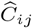, for all values of *C*_*i*_*C*_*j*_.

**Fig. 2.**
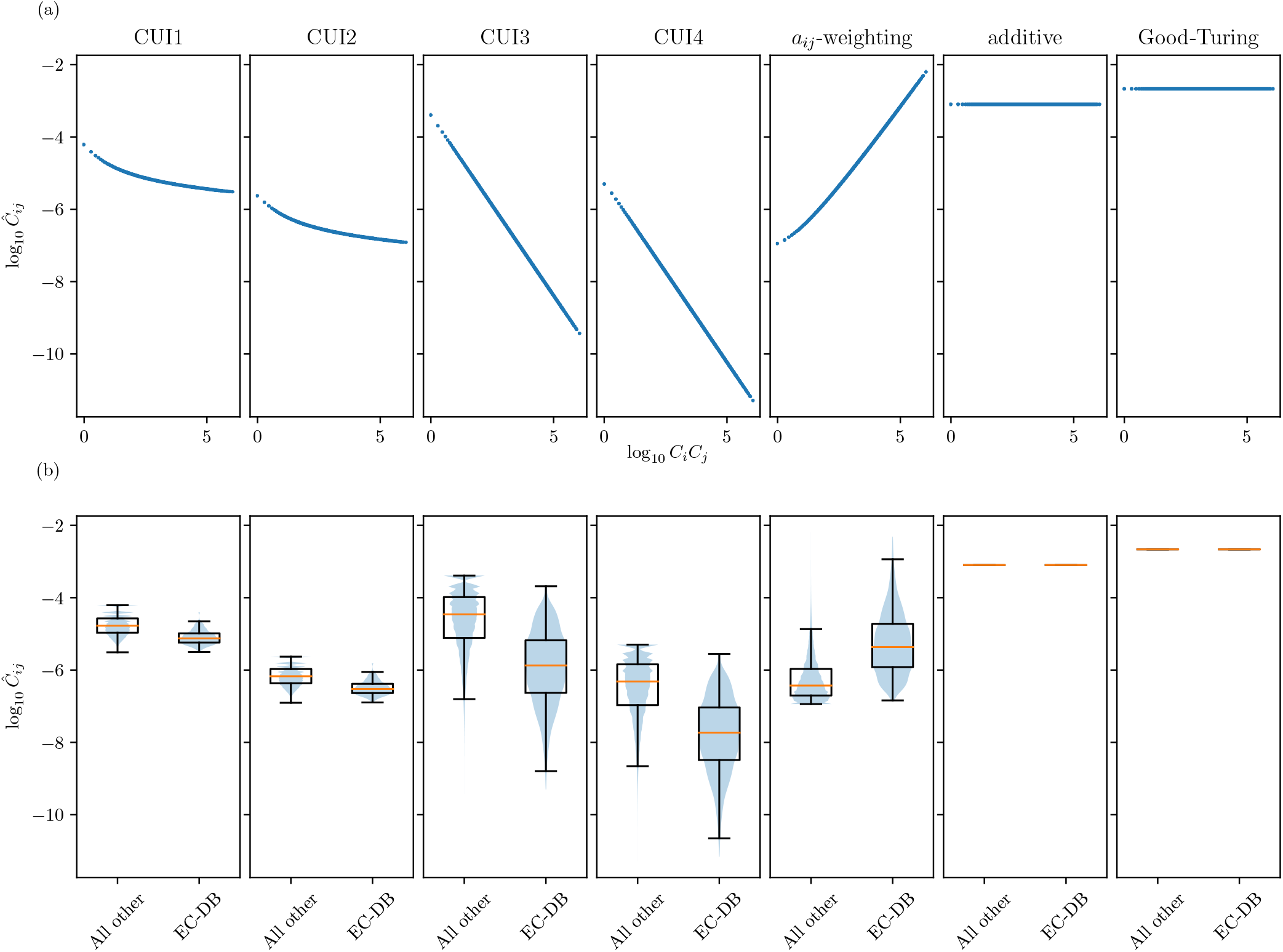
Adjusted bigram counts for unseen bigrams after smoothing on genuine human domain architecture data. For all smoothing methods, we use an e!ective pseudocount of *Ψ* = 0.0008, as calibrated with (10) and (11). (a) Adjusted bigram count 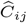 as a function of the product of unigram count *C*_*i*_*C*_*j*_. (b) Distribution of adjusted counts for unseen bigrams containing one extracellular domain and one DNA-binding domain, in either order, (EC-DB) and all other unseen bigrams (All other). For *a*-weighting, *F* (*C*, *C*) = ln(1 + *C C*) and 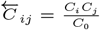 are used here.

We next consider the adjusted counts assigned to our curated set of 5,998 unseen pairs consisting of one DNA-binding and one EC domain (see Methods). Two lines of evidence suggest that these pairs have zero-valued counts because they are incompatible, and not simply because of missing data: First, the products of unigram counts (*C*_*i*_*C*_*j*_) for EC-DB (or DB-EC) pairs are stochastically greater than those for domain pairs in general (*p <* 0.0001, *U* = 3E8, one-sided Mann-Whitney U test), supporting the reliability of these zero-valued observations. In addition, these domains are likely associated with different cellular locations that are not adjacent in the cell.

We compare the distributions of adjusted counts assigned to the 5,998 incompatible pairs and to all other unseen pairs (~ 1E6 bigrams, Figure 2b). As expected, for all four proposed variants in Table 1, the adjusted counts of the incompatible pairs are stochastically smaller than the adjusted counts for all unseen bigrams (Figure 2b, *p <* 0.0001, *U >* 1E9, one-sided Mann-Whitney U test). In contrast, *a*_*ij*_-weighting yields significantly higher adjusted counts for the incompatible bigrams than background (*p <* 0.0001, *U* = 6E9, one-sided Mann-Whitney U test). Additive and Good Turing smoothing do not differentiate between unseen pairs, as can be seen from the horizontal lines.

Note that incompatible subcellular localization is only one of many reasons that a domain pair may be disadvantageous; the “all other” set of unseen bigrams therefore likely contains many additional incompatible pairs that are not members of our curated set.

We have demonstrated that our proposed method respects our assumptions about the reliability of bigrams in multidomain architecture data. We also showed that the alternative *a*_*ij*_-weighting design (20) fails to align with these assumptions. We next assess the ability of each model variant of our design (Table 1) to complete incomplete data.

### Completion of incomplete data

We next assess the ability of each model variant of our design (Table 1) to complete incomplete data. Genuine domain architecture is incomplete for the reasons discussed in the introduction and no gold standard complete data set is available. Instead, we mimic incomplete data by randomly discarding *x*% of domain architectures, *x* ∈ {1%, 2%,…, 10%}, to obtain a reduced set, *R* (*x*), as described in Methods.

We calculate the bigram counts and frequencies in *R*, denoted by 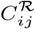 and *P*^*R*^ (*D*_*i*_*D*_*j*_). Using these statistics, we apply smoothing on *C*_*i*_^*R*^_*j*_ to obtain the adjusted bigram counts and frequencies on the incomplete set, denoted by 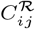 and *P* (*D*_*i*_ *D*_*j*_). For the purpose of these calculations, unigram frequencies are estimated on the complete set *A* to ensure that all domain superfamilies in the original domain alphabet are retained. To account for sample variance, 1000 replicates of this process were conducted for each value of *x*.

We compare the ability of the proposed method variants to complete incomplete data using three figures of merit, described in Methods. The performance of additive smoothing is included as a baseline. For each variant, the smoothing parameter *ε* is selected such that it produces the same relative change in observed bigram count (10) as additive smoothing with pseudocount *Ψ*. This “effective pseudocount” is used as the independent variable. Figure 3 illustrates smoothing performance on incomplete data from which 2% of domain architectures, selected at random, have been removed. Experiments with other values of *x* exhibit the similar trends (Figures S2-S5), with a higher value of *ε* required for optimal performance with increasing incompleteness of the data.

In terms of bigram frequency deviation (Figure 3(a)), all four variants outperform additive smoothing. Among the variants, the smoothing methods based on interpolation to the unigram model (*CUI2* and *CUI4*, solid lines) are less sensitive to the choice of the smoothing parameter than those based on interpolation to the uniform model (Figure 3(b)). Nevertheless, the differences among the variants are small compared to their collective improvement over additive smoothing.

**Fig. 3.**
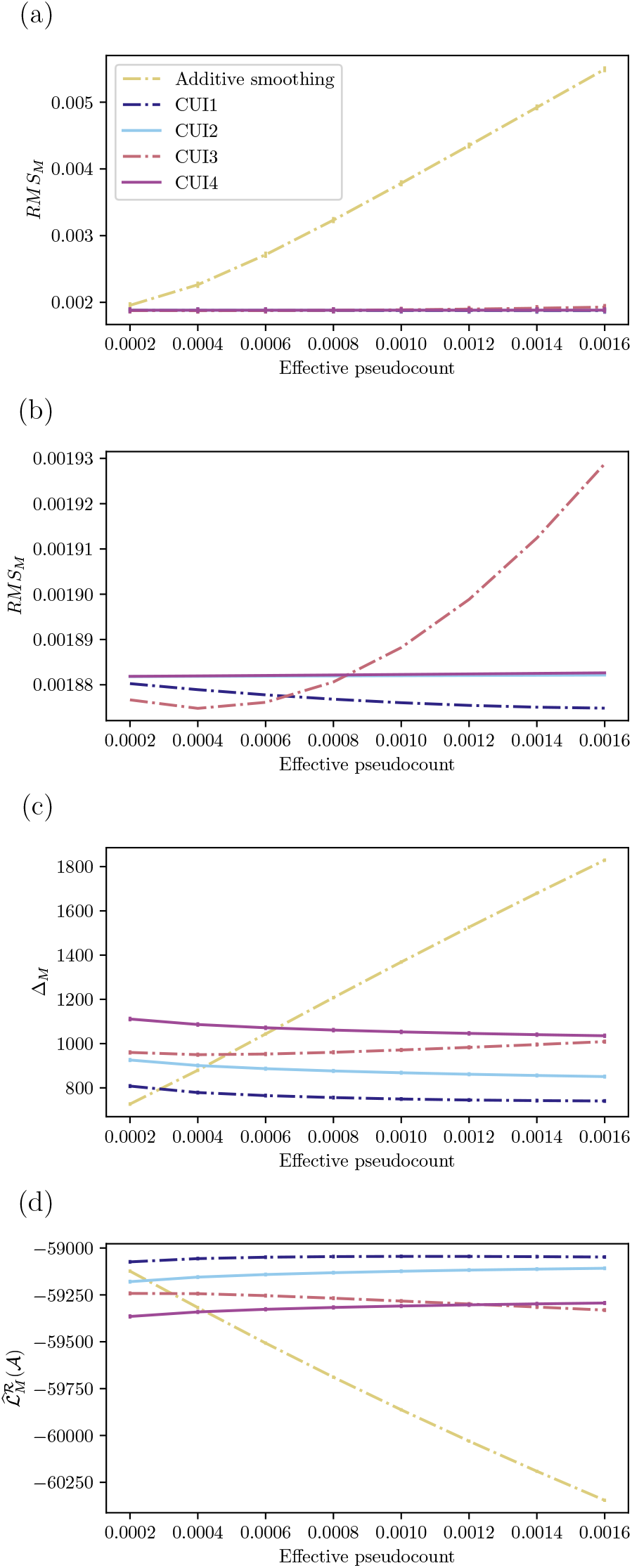
Performance of proposed model variants on smoothing incomplete human domain architecture data, obtained by discarding 2% of all genuine domain architectures, selected at random. Root mean square bigram frequency deviation (a) with detail (b). Architecture probability deviation (c). Log probability of genuine domain architectures (d). All measures are averaged over 1, 000 random experiments. For error bars representing variance, see Figures S2-S5.

For figures of merit based on genuine domain architecture probabilities, either in terms of sum of log of architecture probability deviation (Figure 3(c)) or log probability of the complete genuine set (Figure 3(d)), performance of additive smoothing rapidly worsens as the pseudocount increases. In comparison, all four model variants show stable trends, indicating robustness to the choice of the smoothing parameter *ε*. The variants based on *F* (*C*_*i*_, *C*_*j*_) = ln(1 + *C*_*i*_*C*_*j*_) (*CUI1* and *CUI2*, blue lines) generally perform better than those based on *F* (*C*_*i*_, *C*_*j*_) = *C*_*i*_*C*_*j*_ (*CUI3* and *CUI4*, red lines), which is not surprising given that the values of *C*_*i*_*C*_*j*_ can range over six orders of magnitude. We also observe that interpolation to the uniform distribution (*CUI1* and *CUI3*, dashed lines) is generally better than interpolation to the unigram model (*CUI2* and *CUI4*, solid lines) with respect to these measures, which represent the impact of smoothing on all domain architectures, combined. As we will see in the next section, the relative benefits of the two target distributions are more nuanced when focusing on different classes of domain architectures.

## Preservation of biological characteristics of multidomain architectures

In most eukaryotic genomes, the distribution of domain architectures exhibits properties that deviate substantially from chance expectation and are driven by the genome structure, evolutionary processes, and functional requirements. The adjustment to bigram counts due to smoothing should be su$ciently small so that the smoothed data exhibits the same characteristics as the genuine, unsmoothed data. Here, we probe how well our smoothing method preserves three such properties: domain repetitiveness, domain promiscuity, and domain order.

### Domain Repetitiveness

Tandem arrays of repeated domains and motifs are a ubiquitous feature of proteomes throughout the Tree of Life [13, 25]. The propensity to form tandem arrays varies substantially across domain families. While domain families that tend to form repetitive arrays appear to have independent origins and are structurally and functionally diverse, certain commonalities exist. Many repeat-containing proteins have functions associated with protein or nucleic acid binding, transcriptional regulation, and cell-cell communication and adhesion [13]. In contrast to tandem arrays in non-coding DNA, which exhibit continual expansion and contraction, protein repeats show much greater stability over evolutionary time. A sizable majority of tandem repeats in the human proteome are highly conserved within metazoa, with respect to both sequence and copy number [32].

Given the prevalence and conservation of repetitive domains in human proteins, we seek a smoothing method that preserves the repetitive tendencies of individual domain families and of the genome as a whole. We assess the tendency of a domain to follow another instance in the same domain superfamily using the homogeneous bigram (or *homobigram*) probability, *P* (*D*_*i*_*D*_*i*_). In genuine data, the homobigram probabilities of the 10 most highly repeated domains in primate genomes [12] are significantly elevated compared to homobigram probabilities in the complete set (Figure 4a, *p <* 0.0001,*U* = 53, one-sided Mann-Whitney U test), demonstrating the effectiveness of this measure. Further, when using CUI2 and a moderate value of the smoothing parameter, the homobigram probabilities of genuine pairs and smoothed pairs are strongly correlated (Figure 4(b)), suggesting that our method introduces minimal change to domain repetitiveness.

**Fig. 4.**
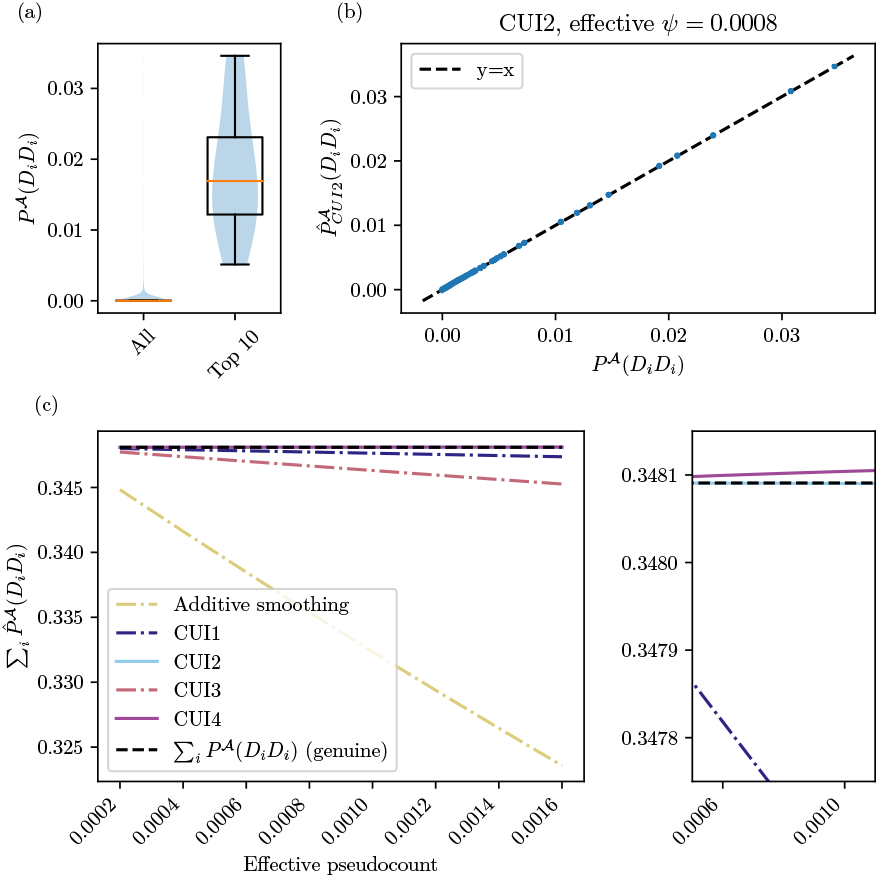
Performance of proposed model variants on recapitulation of domain repetitiveness. (a) Homobigram probability distribution for all domains (All) and for the 10 domains with most copies found in tandem arrays in primate genomes (Top 10): Immunoglobulin, EGF/Laminin, Spectrin repeat, Fibronectin type III, LDL receptor-like module, ARM repeat, Beta-beta-alpha-zinc fingers, Cadherin-like, Complement control module/SRC domain, and Growth factor receptor domains [12]. (b) Smoothed homobigram probabilities plotted against genuine homobigram probabilities. (c) Left: Genome repetitiveness index derived from smoothed data as a function of effective pseudocount. Right: Detailed view.

We next asked whether smoothing continues to preserve repetitiveness as *Ψ* increases and whether other variants exhibit similar behavior. We use the summary statistic ∑_*i*_*P* (*D D*) = ∑_*i*_ *P* (*D*_*i*_)*P* (*D*_*i*_|*D*_*i*_) as a measure of overall genome repetitiveness (Figure 4(c)). With additive smoothing, the repetitive signal degrades substantially, *Ψ* increases. All four CUI variants perform well in comparison, although the variants that interpolate to the uniform model (CUI1, CUI3) result in a greater deviation from the unsmoothed repetitiveness for higher values of *Ψ*. In contrast, the variants that interpolate to the unigram distribution (CUI2, CUI4) almost completely preserve genuine genome repetitiveness (Figure 4(c), right hand panel).

### Domain promiscuity

The *promiscuity* of a domain refers to its propensity to co-occur with other domains [26]. A domain’s promiscuity is often linked to its function. For example, domains that mediate interactions with other molecules (e.g., SH2, SH3, PDZ) are frequently highly promiscuous. These domains combine with a wide variety of catalytic or regulatory domains, giving rise to diverse pathways in which similar catalytic or regulatory functions are applied to different interaction partners [29].

A variety of promiscuity measures have been proposed [38], including the number of distinct direct neighbors

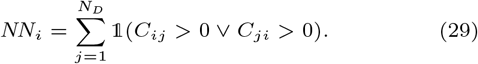

The distribution of *NN* (and many other promiscuity measures) has been shown to follow a power-law [5, 22, 39], indicating that many domains families appear alone or with a small number of other domain families, while a few are found adjacent to many different domains families.

The *NN* measure cannot be used with smoothed data, because with smoothing, every pair of domains has a non-zero bigram probability. Instead, we use entropy, denoted *π*_*i*_, which captures how unevenly the probability of adjacency is distributed among all possible neighbors of a domain *D*_*i*_,

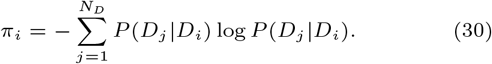

In genuine data, *π*_*i*_ correlates well with log *NN*_*i*_ (Figure 5(a)), suggesting it is a good proxy for *NN*_*i*_. Like *NN*_*i*_, 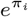 follows a power law as illustrated in Figure 5(c), which was generated as follows: The observed values of 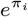 were partitioned into 50 uniformly spaced bins and the log of the number of domains that fall into the each bin was plotted against the corresponding value of *π*. The slope of this log-log scale plot (i.e., the power-law exponent) captures the distribution of domain promiscuities across the genome; a greater absolute value indicates a greater span between the most promiscuous and the least promiscuous domain in the genome. Thus, the power-law exponent serves as a summary statistic representing the genome-wide promiscuity distribution.

**Fig. 5.**
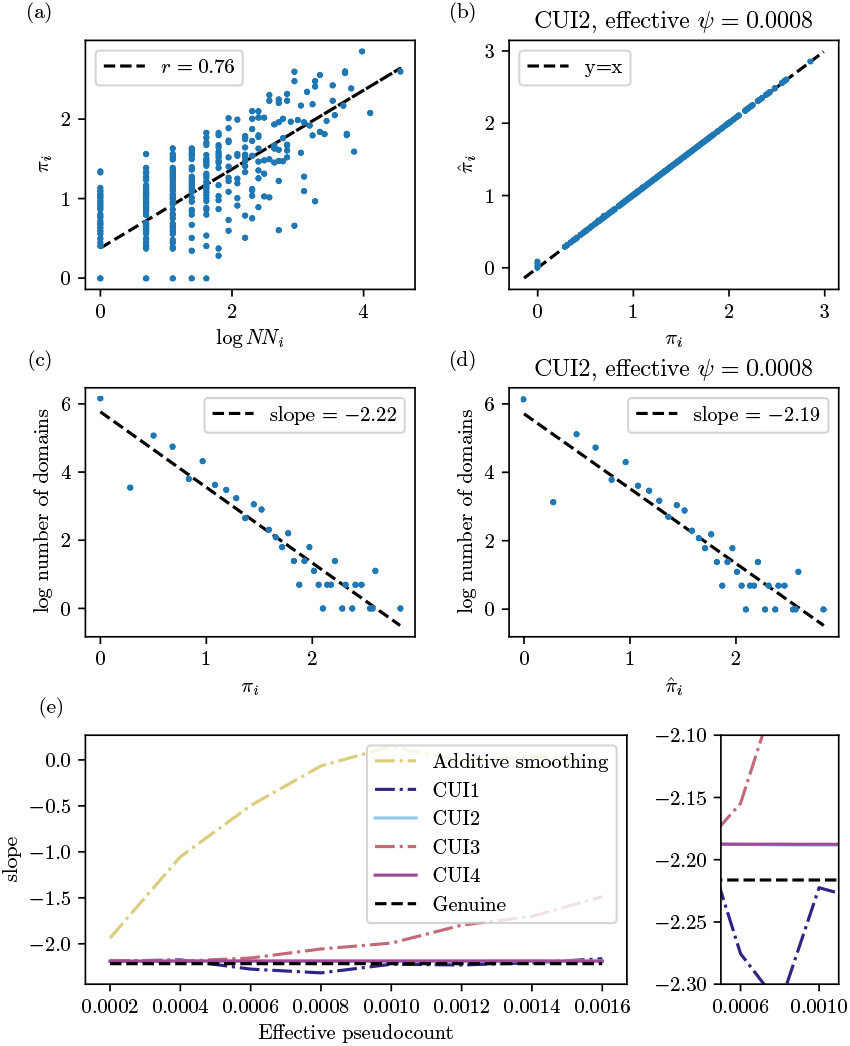
Performance of proposed model variants on recapitulation of domain promiscuity. (a) Entropy as a proxy for number of direct neighbors in genuine data. (b) Entropy derived from smoothed data plotted against genuine entropy. (c) Power-law approximation of distribution of entropy derived from genuine data. (d) Power-law approximation of distribution of entropy derived from smoothed data. (e) Left: Genome-wide promiscuity represented by the slope of the power law derived from smoothed data plotted as a function of effective pseudocount. Right: detailed view.

Here, we probe how well smoothing preserves a domain’s characteristic propensity to co-occur with other domains. Comparison of the entropy of *D*_*i*_ calculated using raw bigram counts and bigram counts smoothed with CUI2 shows minimal change in domain promiscuity for an intermediate value of the smoothing parameter (Figure 5(b)). This is also true of the genome-wide promiscuity distribution: the estimated power-law exponents obtained with genuine and CUI2-smoothed data (−2.22 versus −2.19) differ little when *Ψ* = 0.0008 (Figures 5(c) and 5(d)).

We next compare how well different smoothing variants preserve the genome-wide promiscuity distribution observed in genuine data, as the smoothing parameter increases (Figure 5(e)). With additive smoothing, the variance in genome-wide promiscuity decreases rapidly as *Ψ* increases, diluting the link between a domain’s function and its propensity to partner with diverse domains. In comparison, the impact of CUI-smoothing is modest. For the smallest values of *Ψ*, all four variants have minimal impact on genome wide promiscuity. As *Ψ* increases, we see some deviation from the genuine genome-wide promiscuity distribution when the data is smoothed with CUI1 and more so with CUI3. In contrast, CUI2 and CUI4 are relatively insensitive to increasing *Ψ*. With both of these variants, the smoothed distribution remains close to the genuine distribution even when *Ψ* is large.

For both repetitiveness and promiscuity assessments, among our model variants, interpolation to the unigram model (*CUI2* and *CUI4*) preserves genuine measures more than interpolation to the uniform distribution (*CUI1* and *CUI3*). This is because these two measures both emphasize a non-uniform preference in the tendency of domains to form neighbors, disfavoring interpolation to a uniform distribution of bigram counts.

### Domain order and adjacency

The function of a multidomain protein depends not only on its domain content, but also on the order in which those domains occur. There are many examples of multidomain proteins where three or more domains must appear in a precise order for the protein to function correctly. In multidomain signaling proteins, domain order precisely controls the sequence of conformational changes, post-translational modifications, and binding to interaction partners required to recognize a signal and initiate a response [29]. Scaffold proteins are made up of modular interaction domains that assemble components of a signaling complex at a precise location in the cell, thereby controlling the specificity, timing and dynamics of the downstream signaling pathway [18]. Again, a precise domain order is required to juxtapose the constituents of the signaling complex in the correct spatial configuration and temporal sequence.

In many of these proteins, ordering constraints extend over three or more domains, leading us to consider whether the bigram model is su$cient to capture the constraints on domain order in such proteins. The probability of a domain architecture can be estimated using a first order approximation (Equation 2), where the individual terms in the equation are conditional probabilities estimated from smoothed bigram counts. These terms reflect constraints on domain adjacency, but do not capture longer range constraints. One solution would be to use a higher-order approximation wherein conditional probabilities depend on a prefix of *n* − 1 domains. However, *n*−grams in domain architecture data are too sparse to support *n*−gram models for *n* ≥ 3. For example, in our human data, there are 2, 526 distinct bigrams corresponding 0.2% of all possible bigrams. The number of distinct trigrams is slightly smaller (2, 355), but these correspond to a much smaller percentage (0.00017%) of the number of possible trigrams. As *n* increases, the percentage of *n*−grams that are observed will decrease by a factor of *N* for every increment in *n*.

Noting that a substantial majority of domain pairs are only observed in one of two possible orders [1, 2, 4, 24], we posit that bigrams implicitly encode longer range constraints on domain order in multidomain architectures. This suggests that a first-order approximation based on bigram frequencies might offer a reasonable approximation of domain order.

To probe this hypothesis empirically, we compare the frequency of each trigram observed in human multidomain architectures, *P* (*D*_*i*_*D*_*j*_ *D*_*k*_), with its first order approximation, *P* (*D*_*i*_)*P* (*D*_*j*_ |*D*_*i*_)*P* (*D*_*k*_|*D*_*j*_) in Figure 6. The first order approximation closely matches the observed trigram frequencies for frequently observed trigrams; for less frequently observed trigrams it is less accurate, but the deviations remain within the expected sampling error of three standard deviations.

**Fig. 6.**
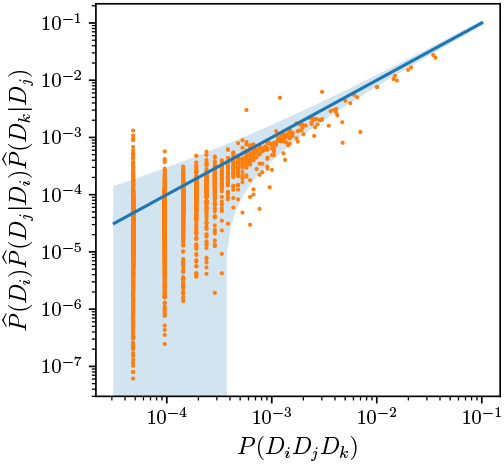
Observed trigram probabilities versus the first order estimation obtained using empirical bigram probabilities (orange dots). The main diagonal (blue solid line) represents exact agreement of the observed and estimated trigram probabilities. The shaded blue envelope shows the 3 *σ* region, where 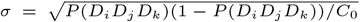 is the standard deviation obtained when estimating the trigram probability from *C*_0_ independent trials.

The results in Figure 6 suggest that bigram frequencies may be suffciently constrained to capture longer range order constraints. To explore this hypothesis for concrete cases, we examined the performance of the first order approximation on five specific examples of signaling and scaffolding proteins (Figure 7, left), in which domain order is likely crucial to function. Two of these, CASK and DLG 1, are scaffold proteins that organize signaling complexes at cell junctions [8]. In neuronal cells, they mediate synaptic protein complex formation and regulation [15].

**Fig. 7.**
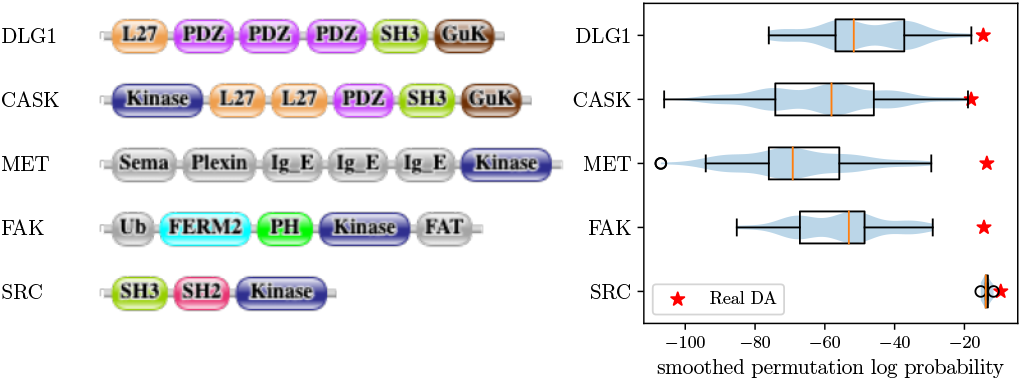
Left: The domain architectures of five proteins, wherein domain order is important for protein function (see text). Domain architecture schematics generated with domain-gfx (https://github.com/EBI-Metagenomics/domain-gfx). Right: The probabilities of all domain architectures obtained by permuting the order of the domains in the architectures shown at right. Architecture probabilities calculated using the first order approximation (Equation 2), with CUI2-smoothed bigram frequencies (*Ψ* = 0.0008). In each case, the probability of the genuine architecture is indicated by a red star.

The remaining three examples are members of the tyrosine kinase family of multidomain signaling proteins. The Focal Adhesion Kinase 1 (FAK) is a receptor that performs numerous signaling functions at the interface between cell-matrix adhesions and the cytoskeleton [31]. FAK integrates multiple molecular stimuli via its five domains, which mediate different types of molecular interactions, including phospholipid binding, localization to the plasma membrane, phosphorylation and dimerization. The spatial and temporal sequencing of these interactions depend on the order of FAK’s constituent domains. The MET receptor tyrosine kinase orchestrates an extensive network of cell surface proteins through interactions with an astounding number of molecules, including other RTKs, co-receptors, adhesion molecules, phosphatases, and other types of receptors [16]. Domain order in both FAK and MET is likely strictly conserved by pleiotropic constraints.

The final example, SRC, is a non-receptor tyrosine kinase with a diverse interaction network, including FAK and MET. We selected SRC as a potentially challenging case for the bigram model. It has only three domains (SH3-SH2-Kinase), in contrast to the other four examples, which have five or more. Moreover, SH2 and SH3 are among the few domain pairs that are observed in both orientations [24].

For each of these examples, we calculated the probabilities of the genuine domain architecture and of all permutations of its constituent domains using CUI2-smoothed bigram frequencies. (Note that calculating the probabilities of permutations is only possible with smoothed bigram data, because many permutations contain unseen bigrams.) If the first order approximation were su$cient to capture domain content, but not domain order, we would expect the estimated probability of the genuine domain architecture to be similar to the estimated probabilities of its permutations. If, in contrast, the first order approximation calculated with smoothed bigram frequencies does reflect domain order over longer ranges, probability of the genuine domain architecture should be higher than the probabilities of its permutations. This is, in fact, what we observe (Figure 7, right). For all five examples, including SRC, the genuine architecture has the highest probability.

## Discussion

We have introduced a customizable unified interpolation (CUI) smoothing framework for *n*-gram models, in which the trust parameters *a* and *b* and the target *n*-gram count distribution 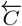 can be chosen to suit the characteristics of the data and the desired smoothing behavior.

Building on this framework, we designed a suite of bigram smoothing methods for incomplete multidomain architecture data. We focus on bigram smoothing because the sparsity of multidomain architecture precludes the use of higher order *n*-gram models. Our methods are tailored to the reliability assumptions appropriate for our application. Specifically, this design rests on the assumption that bigram counts involving domains with high unigram counts are more reliable, and is designed to make smaller adjustments to such counts. We also posit that some unseen bigrams are not observed because the constituent domains are biologically incompatible, while others are zero-valued due to data incompleteness. A consequence of these assumptions is that unseen pairs composed of high-frequency domains are treated as truly incompatible and assigned very low counts. Unseen pairs of low-frequency domains may be equally incompatible, but the data do not provide the corroborating evidence needed to distinguish them from undersampled pairs.

We demonstrated, both theoretically and empirically, that these methods (1) assign non-zero counts to unseen pairs while distinguishing truly incompatible pairs from those that are merely undersampled, (2) adjust the counts of seen pairs to account for incompleteness, and (3) preserve the characteristic properties of multidomain architecture data, including repetitiveness, promiscuity, and order constraints.

The strength of the CUI framework lies in its flexibility: the trust parameters and target distribution can be chosen to encode different assumptions about the data. This flexibility also makes explicit that the assumptions appropriate for multidomain architectures differ from those underlying NLP. Parameter choices that are natural for language corpora are readily expressed within our framework, but yield smoothing behavior poorly suited to multidomain data. Our specific design instead encodes the assumptions appropriate for multidomain architecture data.

Several directions remain open for further work. The differing behavior of model variants across domain classes motivates a design in which distinct target distributions are applied to seen and unseen bigrams. Beyond the three characteristics examined here, additional features of multidomain proteins are exciting directions for further analysis, including the domain architecture length distribution, asymmetry in domain order, and domains that rarely or never combine with other domain families. Different choices of the trust parameters and target distribution can tailor the framework to other types of bigram data; applying the framework to other corpora is a promising direction to explore.

## Data availability

Implementations of the smoothing methods, the scripts used to generate all results presented in this paper, and the curated lists of extracellular and DNA-binding domains are available at https://codeberg.org/xcui297/protein-domain-smoothing.

## Competing interests

No competing interest is declared.

Author contributions statement

The domain bigram smoothing problem was proposed by XC and DD; the mathematical models were developed by XC and GI, with input from DD; XC conceived and conducted experiment(s), with input from GI and DD, and analyzed the results; all authors wrote and reviewed the manuscript.

## Acknowledgments

The authors thank Dr. Maureen Stolzer for discussions on modeling multidomain architecture data. This work is supported in part by funds from the National Science Foundation (NSF Grants DBI-1838344 and DBI-1759943 to DD and # 2342349 to GI) and the Center for Nonlinear Analysis.

## Footnotes

1 Notice that when the trust parameters *a*_*ij*_ = *b*_*ij*_ = 1, and the target pairwise count distribution 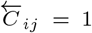, then this is equivalent to the standard additive smoothing (5) as both result in distributions with the same adjusted bigram probabilities.

## Notes

https://codeberg.org/xcui297/protein-domain-smoothing

### Competing Interest Statement

The authors have declared no competing interest.

