## Supplementary Information for "A unified smoothing framework for protein domain bigram model"

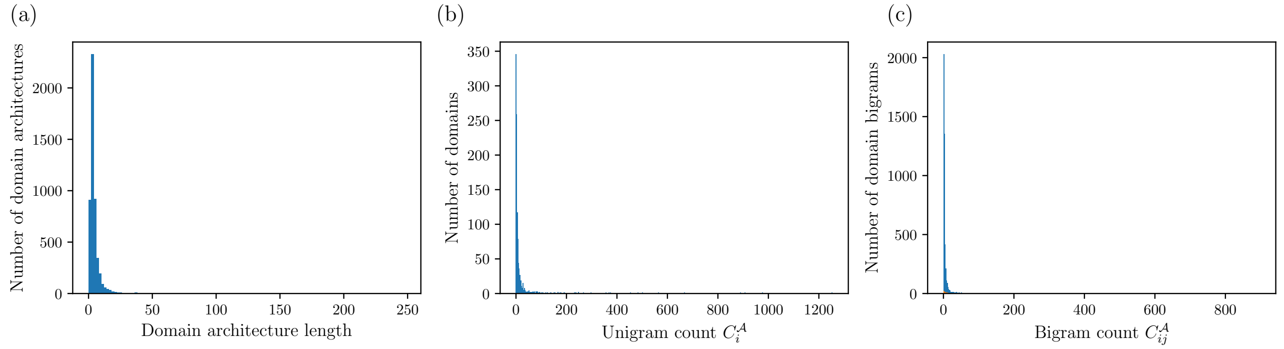


**Fig. S1.** Full distributions of (a) domain architecture length, (b) unigram counts, and (c) bigram count.

**Table S1.** Effective pseudocount.

| $\psi$ | 0.0002000 | 0.0004000 | 0.0006000 | 0.0008000 | 0.0010000 | 0.0012000 | 0.0014000 | 0.0016000 |
|---|---|---|---|---|---|---|---|---|
| $\varepsilon^*_{CUI1}(\psi)$ | 0.0000108 | 0.0000216 | 0.0000324 | 0.0000431 | 0.0000539 | 0.0000647 | 0.0000755 | 0.0000863 |
| $\varepsilon^*_{CUI2}(\psi)$ | 0.0000004 | 0.0000009 | 0.0000013 | 0.0000017 | 0.0000022 | 0.0000026 | 0.0000030 | 0.0000035 |
| $\varepsilon^*_{CUI3}(\psi)$ | 0.0001043 | 0.0002088 | 0.0003133 | 0.0004180 | 0.0005228 | 0.0006277 | 0.0007328 | 0.0008379 |
| $\varepsilon^*_{CUI4}(\psi)$ | 0.0000014 | 0.0000029 | 0.0000043 | 0.0000058 | 0.0000073 | 0.0000087 | 0.0000102 | 0.0000117 |

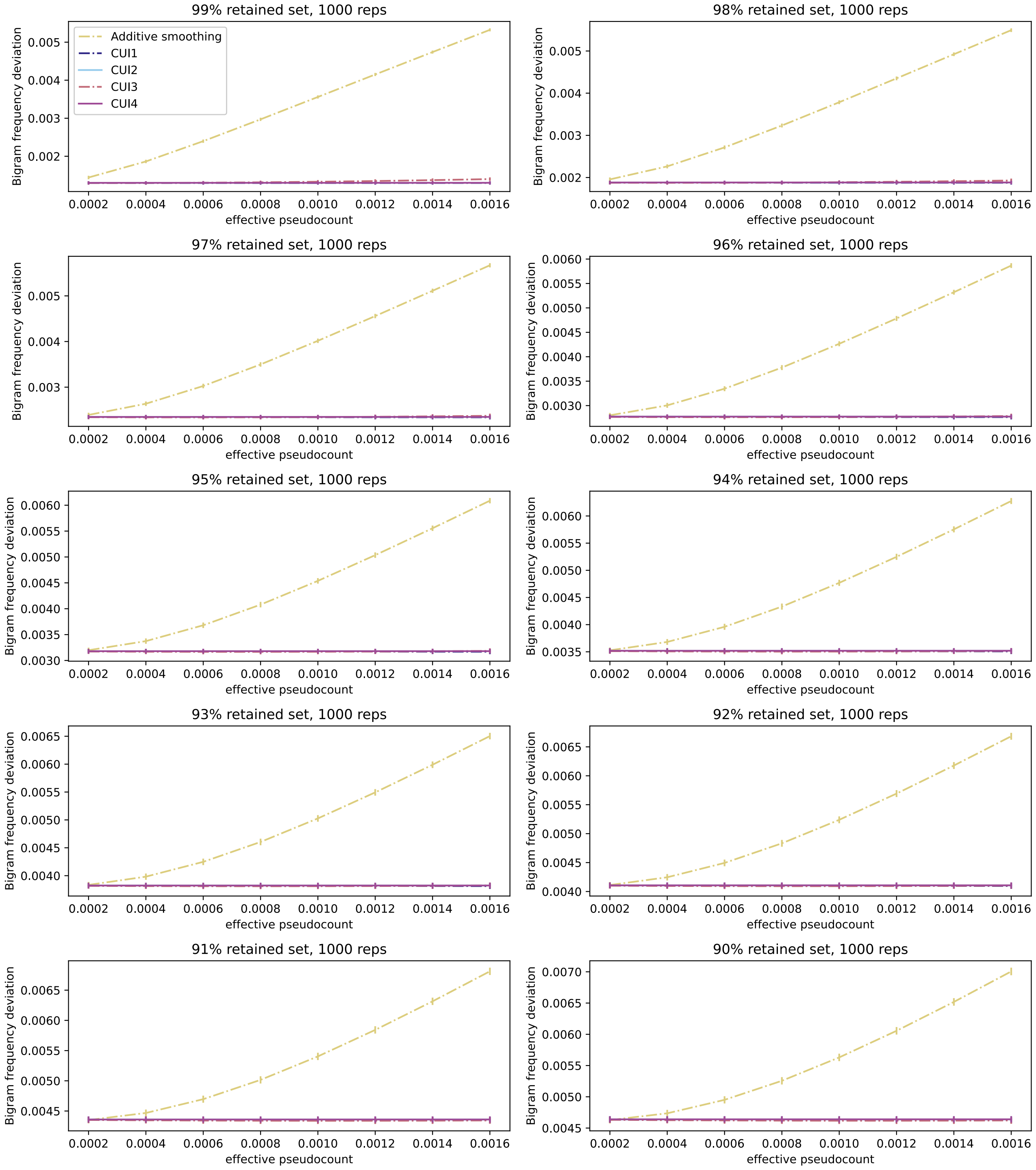


**Fig. S2.** Performance of proposed model variants on smoothing incomplete genuine human domain architecture data, assessed with root mean square bigram frequency deviation. Incomplete data sets were constructed by randomly discarding $x\%$ of domain architectures, $x \in \{1\%, 2\%, \ldots, 10\%\}$. All measures are averaged over $1,000$ randomly generated reduced sets. Error bars show standard error across the $1,000$ replicates.

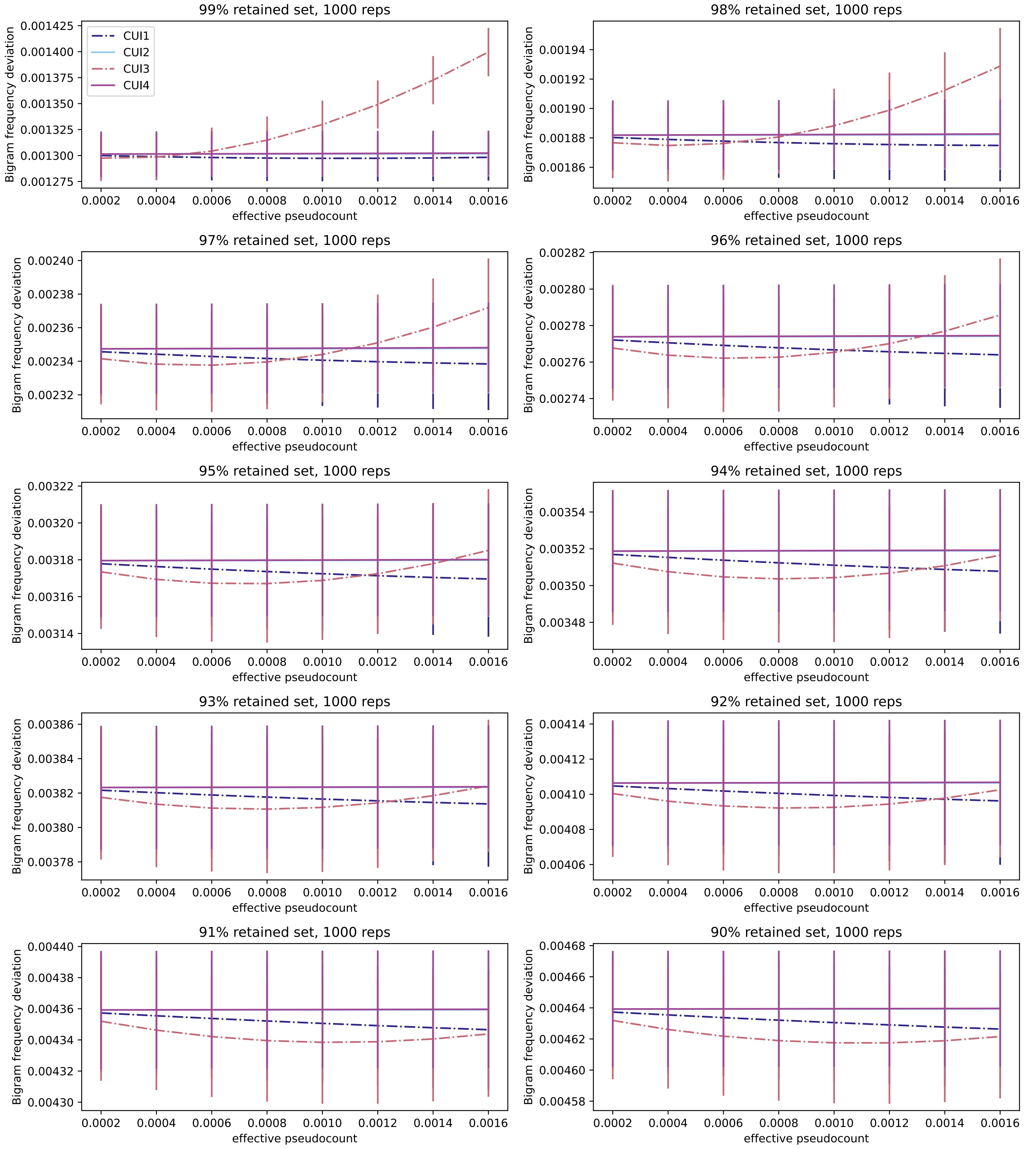


**Fig. S3.** Detailed view of performance of proposed model variants on smoothing incomplete genuine human domain architecture data, assessed with root mean square bigram frequency deviation. Incomplete data sets were constructed by randomly discarding $x\%$ of domain architectures, $x \in \{1\%, 2\%, \ldots, 10\%\}$. All measures are averaged over $1,000$ randomly generated reduced sets. Error bars show standard error across the $1,000$ replicates.

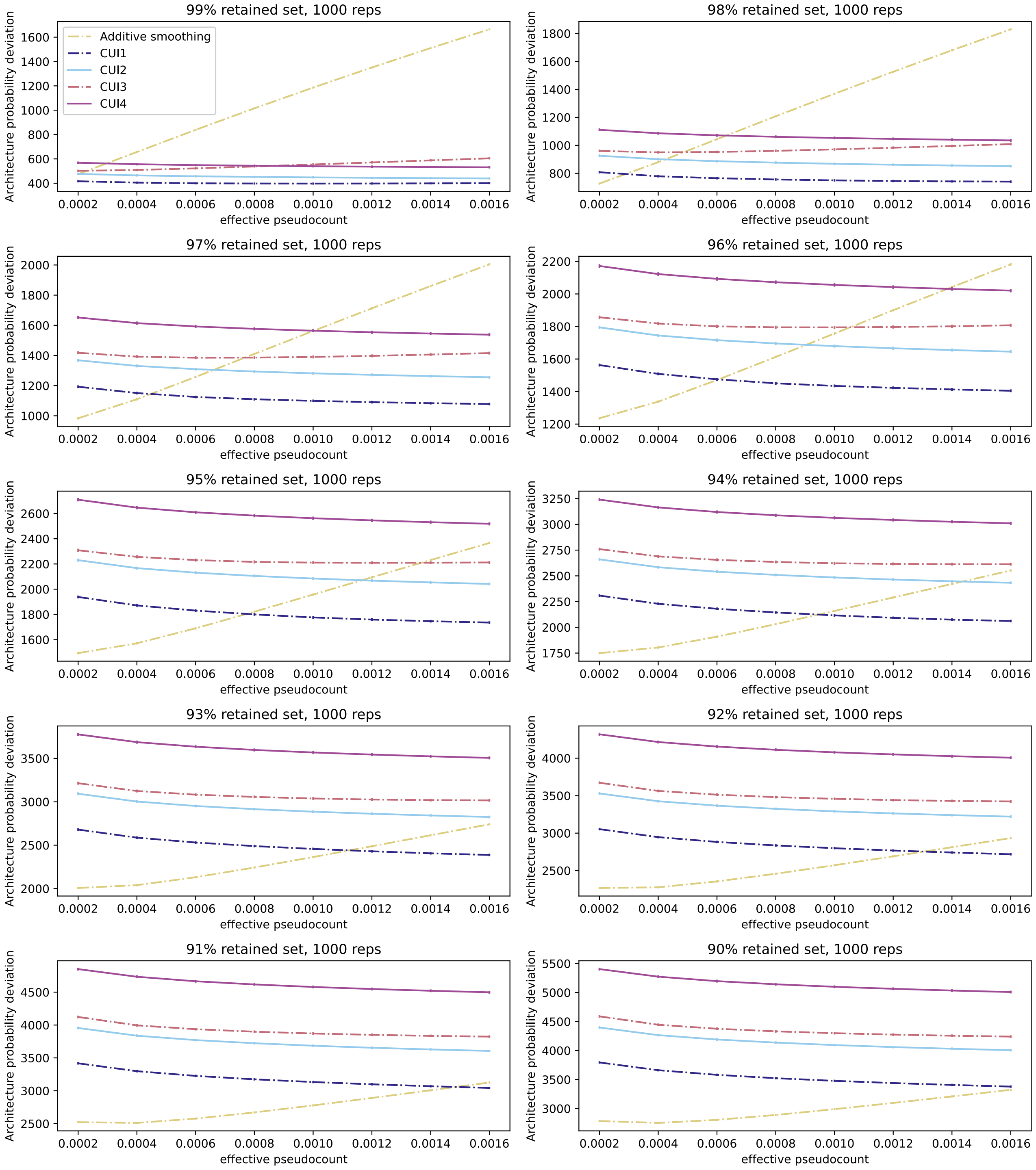


**Fig. S4.** Performance of proposed model variants on smoothing incomplete genuine human domain architecture data, assessed with architecture probability deviation. Incomplete data sets were constructed by randomly discarding $x\%$ of domain architectures, $x \in \{1\%, 2\%, \ldots, 10\%\}$. All measures are averaged over $1,000$ randomly generated reduced sets. Error bars show standard error across the $1,000$ replicates.

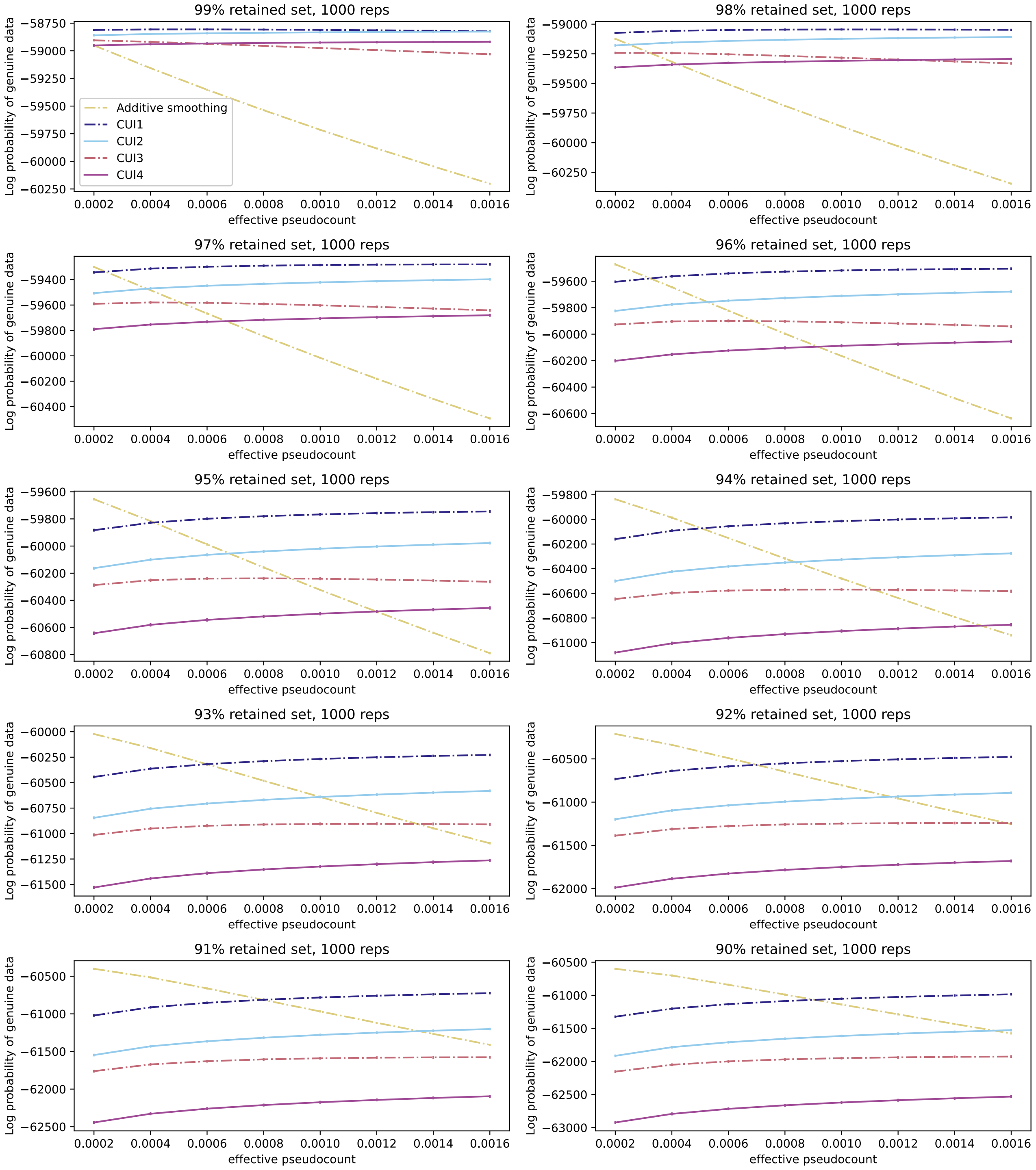


**Fig. S5.** Performance of proposed model variants on smoothing incomplete genuine human domain architecture data, assessed with log probability of genuine domain architectures. Incomplete data sets were constructed by randomly discarding $x\%$ of domain architectures, $x \in \{1\%, 2\%, \ldots, 10\%\}$. All measures are averaged over $1,000$ randomly generated reduced sets. Error bars show standard error across the $1,000$ replicates.

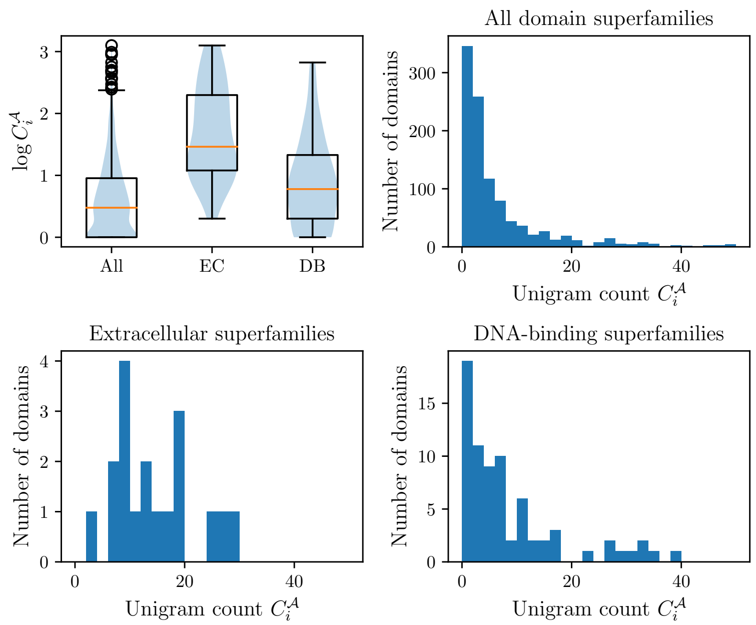


**Fig. S6.** Distributions of unigram counts of all domain superfamilies, extracellular (EC) superfamilies, and DNA-binding (DB) superfamilies. Both EC and DB domains have unigram counts stochastically greater than those of domain superfamilies overall ($p < 0.0001$, $U > 4E3$, one-sided Mann-Whitney U test). The products of unigram counts ($C_i C_j$) for EC-DB (or DB-EC) pairs are also stochastically larger than those for domain pairs in general ($p < 0.0001$, $U = 3E8$, one-sided Mann-Whitney U test)

**Table S2.** List of extracellular domains.

| SUPFAM ID | Domain superfamily name |
|---|---|
| 100895 | Kazal-type serine protease inhibitors |
| 49265 | Fibronectin type III |
| 49854 | Spermadhesin, CUB domain |
| 49899 | Concanavalin A-like lectins/glucanases |
| 56436 | C-type lectin-like |
| 57184 | Growth factor receptor domain |
| 57196 | EGF/Laminin |
| 57440 | Kringle-like |
| 57535 | Complement control module/SCR domain |
| 57603 | FnI-like domain |
| 57610 | Thyroglobulin type-1 domain |
| 57630 | GLA-domain |
| 69848 | LCCL domain |
| 82671 | SEA domain |
| 82895 | TSP-1 type 1 repeat |
| 90188 | Somatomedin B domain |
| 101912 | Sema domain |
| 103575 | Plexin repeat |
| 47686 | Anaphylotoxins (complement system) |
| 47874 | Annexin |
| 50242 | TIMP-like |
| 50923 | Hemopexin-like domain |
| 52058 | L domain-like |
| 54511 | GFP-like |
| 56496 | Fibrinogen C-terminal domain-like |
| 82153 | FAS1 domain |
| 48726 | Immunoglobulin |
| 57424 | LDL receptor-like module |
| 56487 | SRCR-like |
| 57362 | BPTI-like |
| 63501 | Frizzled cysteine-rich domain |
| 57256 | Elafin-like |
| 57414 | Hairpin loop containing domain-like |
| 57492 | Trefoil |
| 57302 | Snake toxin-like |

**Table S3.** List of DNA-binding domains.

| SUPFAM ID | Domain superfamily | SUPFAM ID | Domain superfamily |
|---|---|---|---|
| 56281 | Metallo-hydrolase/oxidoreductase | 47459 | HLH, helix-loop-helix DNA-binding domain |
| 54791 | Eukaryotic type KH-domain (KH-domain type I) | 46689 | Homeodomain-like |
| 51735 | NAD(P)-binding Rossmann-fold domains | 54076 | RNase A-like |
| 53098 | Ribonuclease H-like | 54928 | RNA-binding domain, RBD |
| 56672 | DNA/RNA polymerases | 50494 | Trypsin-like serine proteases |
| 57667 | beta-beta-alpha zinc fingers | 57756 | Retrovirus zinc finger-like domains |
| 47113 | Histone-fold | 88946 | Sigma2 domain of RNA polymerase sigma factors |
| 53335 | S-adenosyl-L-methionine-dependent methyltransferases | 47413 | lambda repressor-like DNA-binding domains |
| 57959 | Leucine zipper domain | 57701 | Zn2/Cys6 DNA-binding domain |
| 46785 | "Winged helix" DNA-binding domain | 46894 | C-terminal effector domain of the bipartite response regulators |
| 47729 | IHF-like DNA-binding proteins | 53041 | Resolvase-like |
| 55811 | Nudix | 50249 | Nucleic acid-binding proteins |
| 55455 | SRF-like | 46955 | Putative DNA-binding domain |
| 47954 | Cyclin-like | 47095 | HMG-box |
| 54957 | Viral DNA-binding domain | 48452 | TPR-like |
| 49417 | p53-like transcription factors | 56349 | DNA breaking-rejoining enzymes |
| 48150 | DNA-glycosylase | 88723 | PIN domain-like |
| 52425 | Cryptochrome/photolyase, N-terminal domain | 55608 | Homing endonucleases |
| 46767 | Methylated DNA-protein cysteine methyltransferase, C-terminal domain | 56712 | Prokaryotic type I DNA topoisomerase |
| 54197 | HIT-like | 51658 | Xylose isomerase-like |
| 55271 | DNA repair protein MutS, domain I | 143422 | Transposase IS200-like |
| 81296 | E set domains | 81301 | Nucleotidyltransferase |
| 46919 | N-terminal Zn binding domain of HIV integrase | 88697 | PUA domain-like |
| 50486 | FMT C-terminal domain-like | 54160 | Chromo domain-like |
| 82679 | N-utilization substance G protein NusG, N-terminal domain | 75553 | Smc hinge domain |
| 56741 | Eukaryotic DNA topoisomerase I, N-terminal DNA-binding fragment | 54060 | His-Me finger endonucleases |
| 50447 | Translation proteins | 56366 | SMAD MH1 domain |
| 52141 | Uracil-DNA glycosylase-like | 56219 | DNase I-like |
| 48173 | Cryptochrome/photolyase FAD-binding domain | 47598 | Ribbon-helix-helix |
| 53036 | Eukaryotic RPB5 N-terminal domain | 89447 | AbrB/MazE/MraZ-like |
| 88659 | Sigma3 and sigma4 domains of RNA polymerase sigma factors | 46946 | S13-like H2TH domain |
| 52540 | P-loop containing nucleoside triphosphate hydrolases | 52980 | Restriction endonuclease-like |
| 53901 | Thiolase-like | 48371 | ARM repeat |
| 110217 | DNA-binding protein LAG-1 (CSL) | 56300 | Metallo-dependent phosphatases |
| 47802 | DNA polymerase beta, N-terminal domain-like | 56719 | Type II DNA topoisomerase |
| 54171 | DNA-binding domain | 47789 | C-terminal domain of RNA polymerase alpha subunit |
| 63763 | SAND domain-like | 48019 | post-AAA+ oligomerization domain-like |
| 47794 | Rad51 N-terminal domain-like | 47798 | Barrier-to-autointegration factor, BAF |
| 90073 | GCM domain | 81624 | N-terminal domain of MutM-like DNA repair proteins |
| 47819 | HRDC-like | 48295 | TrpR-like |
| 47807 | 5' to 3' exonuclease, C-terminal subdomain | 46950 | Double-stranded DNA-binding domain |
| 102645 | CoaB-like | 56726 | DNA topoisomerase IV, alpha subunit |
| 47454 | A DNA-binding domain in eukaryotic transcription factors | 82927 | Cysteine-rich DNA binding domain, (DM domain) |
| 50809 | XRCC4, N-terminal domain | 47781 | RuvA domain 2-like |
| 74784 | Translin | 54211 | Ribosomal protein S5 domain 2-like |
| 50723 | Core binding factor beta, CBF | 55945 | TATA-box binding protein-like |
| 101224 | HAND domain of the nucleosome remodeling ATPase ISWI | 46774 | ARID-like |
| 56019 | The spindle assembly checkpoint protein mad2 | 54447 | ssDNA-binding transcriptional regulator domain |
| 50978 | WD40 repeat-like | 57783 | Zinc beta-ribbon |
| 117773 | GTF2I-like repeat | | |
